# The extracellular matrix controls stem cell specification and crypt morphology in the developing and adult gut

**DOI:** 10.1101/2021.04.14.439776

**Authors:** R. Ramadan, SM. van Neerven, VM. Wouters, T. Martins Garcia, V. Muncan, OD. Franklin, M. Battle, KS. Carlson, J. Leach, OJ. Sansom, L. Vermeulen, JP. Medema, DJ. Huels

**Affiliations:** Laboratory for Experimental Oncology and Radiobiology, Center for Experimental and Molecular Medicine, Cancer Center Amsterdam, University of Amsterdam, Meibergdreef 9, 1105 AZ, Amsterdam, The Netherlands; Oncode Institute, Meibergdreef 9, 1105 AZ, Amsterdam, The Netherlands; Department of Gastroenterology and Hepatology, Tytgat Institute for Intestinal and Liver Research, Amsterdam Gastroenterology Endocrinology and Metabolism, Amsterdam UMC University of Amsterdam, Amsterdam, The Netherlands; The Medical College of Wisconsin, Department of Cell Biology, Neurobiology, and Anatomy, Milwaukee, Wisconsin, USA; The Blood Research Institute of Wisconsin, part of Versiti, and the Medical College of Wisconsin, Department of Internal Medicine, Milwaukee, Wisconsin, USA; Cancer Research UK Beatson Institute, Garscube Estate, Switchback Road, Glasgow, G61 1BD, UK; Institute of Cancer Sciences, University of Glasgow, Garscube Estate, Switchback Road, G61 1QH, UK

**Keywords:** fetal intestine, extracellular matrix, intestinal stem cells

## Abstract

The rapid renewal of the epithelial gut lining is fuelled by stem cells that reside at the base of intestinal crypts. In recent years, the signal transduction pathways and morphogens that regulate intestinal stem cell self-renewal and differentiation have been extensively characterised. In contrast, although extracellular matrix (ECM) components form an integral part of the intestinal stem cell niche, their direct influence on the cellular composition is less well understood. Here, we set out to systematically compare the effect of two major ECM classes, fibrillar collagen type I and the basement membrane, on the intestinal epithelium. We found that both collagen I and laminin-containing cultures allow growth of small intestinal epithelial cells with all differentiated and undifferentiated cell types present in both cultures, albeit at different ratios. Specific to the collagen culture was a subset of cells with a fetal-like gene expression program. In contrast, laminin, but not collagen IV, increased Lgr5+ stem cells and Paneth cells, and induced crypt-like morphology changes. The transition from a collagen culture to a laminin culture, resembles the gut development *in vivo*. Here, the ECM is dramatically remodelled by mesenchymal cells, which is accompanied by a specific and local expression of the laminin receptor ITGA6 in the crypt-forming epithelium. This laminin:ITGA6 signalling is essential for the stem cell induction and crypt formation *in vitro*. Importantly, deletion of laminin in the adult mouse results in a fetal-like epithelium with a marked reduction of adult intestinal stem cells. Overall, our data support the hypothesis that the formation of intestinal crypts is induced by an increased laminin concentration in the ECM.

## Introduction

The small intestinal epithelium is a rapidly dividing tissue that renews itself from adult intestinal stem cells which reside at the bottom of crypts. The balance between proliferation and differentiation is finely controlled by growth factors and morphogenic signals, creating a niche at the bottom of the crypts that supports stem cell growth and suppresses differentiation (1). The development of the *in vitro* organoid system brought mesenchymal growth factors into focus that are essential for the growth of intestinal stem cells *in vitro* and were then shown to have a similar role *in vivo*, e.g. R-spondin (2– 4). Another requirement of the organoid system is an ECM-derived hydrogel (Matrigel or basement-membrane extracts), providing the required mechanical and signalling cues of the organoid surrounding. These matrices are derived from a mouse sarcoma, the Engelbreth-Holm-Swarm (EHS) tumor, and are rich in laminin, collagen IV and entactin.

The ECM can be divided into two main types, the interstitial matrix surrounding the tissue, and the basement membrane, a thin ECM layer separating the epithelium from the interstitium. The role of the ECM goes beyond just providing a physical scaffold for tissue. Its dynamic remodelling is essential for organ development and in disease (5). For example, it is long established that changes in the ECM have a fundamental role for the development of the mammary gland. Here, collagen I induces ductal branching and in turn causes a remodelling of the basement membrane which is responsible for the differentiation of the mammary gland (6, 7). This ECM remodelling highlights different but coordinated roles for collagen I and the basement membrane in the development and formation of the mammary gland. This raises the possibility that, in the intestine, collagen and the basement membrane, fulfill similar roles during the development as well as maintenance of the intestinal niche. Early studies already observed a fundamental remodelling of the intestinal ECM during gut development (8, 9), however its direct functional relevance on the epithelium is not well described.

In addition to laminin-rich hydrogel cultures (e.g. Matrigel), recent reports showed that intestinal epithelial cells can be maintained in a pure culture of fibrillar collagen type I, with the presence of stem cells as well as differentiated cells(10, 11), suggesting that supplemented basement-membrane components are not essential for the maintenance of intestinal stem cells. However, a recent study by Yui et al. showed that collagen type I induces a reprogramming of the adult intestine to a more fetal-like epithelium with a suppression of intestinal stem cell genes (12).

This ECM remodelling becomes evident in colonic inflammation and regeneration where an increased expression and deposition of collagen type I (14, 15) and a reduction in laminin protein (16) can be detected.

In this study we compared in detail the influence of two different natural ECM derived matrices with similar physical properties, the basement membrane (Matrigel, laminin or collagen IV) and the interstitial matrix (collagen I) on the intestinal epithelium. Both ECMs allowed growth of intestinal cells *in vitro* with all major cell types present in both cultures, however at different ratios. Collagen cultures induced a fetal-like expression program in a subset of cells and contained a reduced number of stem cells as well as Paneth cells. In contrast, laminin signalling (via ITGA6) was responsible for an increase in intestinal stem cells and induction of morphological crypt-like structures. These observations showed a striking similarity to the situation *in vivo*, where laminin deposition by the mesenchyme was enhanced during crypt formation and deletion of laminin in adult mice resulted in stem cell loss and upregulation of fetal-like genes. These data support the idea that the deposition of laminin to the ECM by mesenchymal cells controls intestinal crypt development as well as maintenance *in vitro* and *in vivo*.

## Results

### The ECM controls expression of fetal-like genes and stem cells

To study the impact of the ECM on the small intestinal epithelium we plated organoids in Matrigel and collagen type I. Matrigel is a basement membrane extract that mainly consists of laminin, collagen IV and entactin. Collagen type I is the main component of the interstitium and is the most abundant protein in mammals. We purified small intestinal crypts from adult wildtype mice and grew them as organoids as previously described (2). After establishment, we plated organoids on a pure layer of collagen I (10, 11) or in Matrigel **(Fig. 1a)**. As the growth of intestinal cells embedded in collagen (3D) is only possible with addition of Wnt3a (12), we focused on the 2D model, where cells are plated on a thick layer of collagen I and can be maintained in the same medium as Matrigel cultures.

**Fig. 1.**
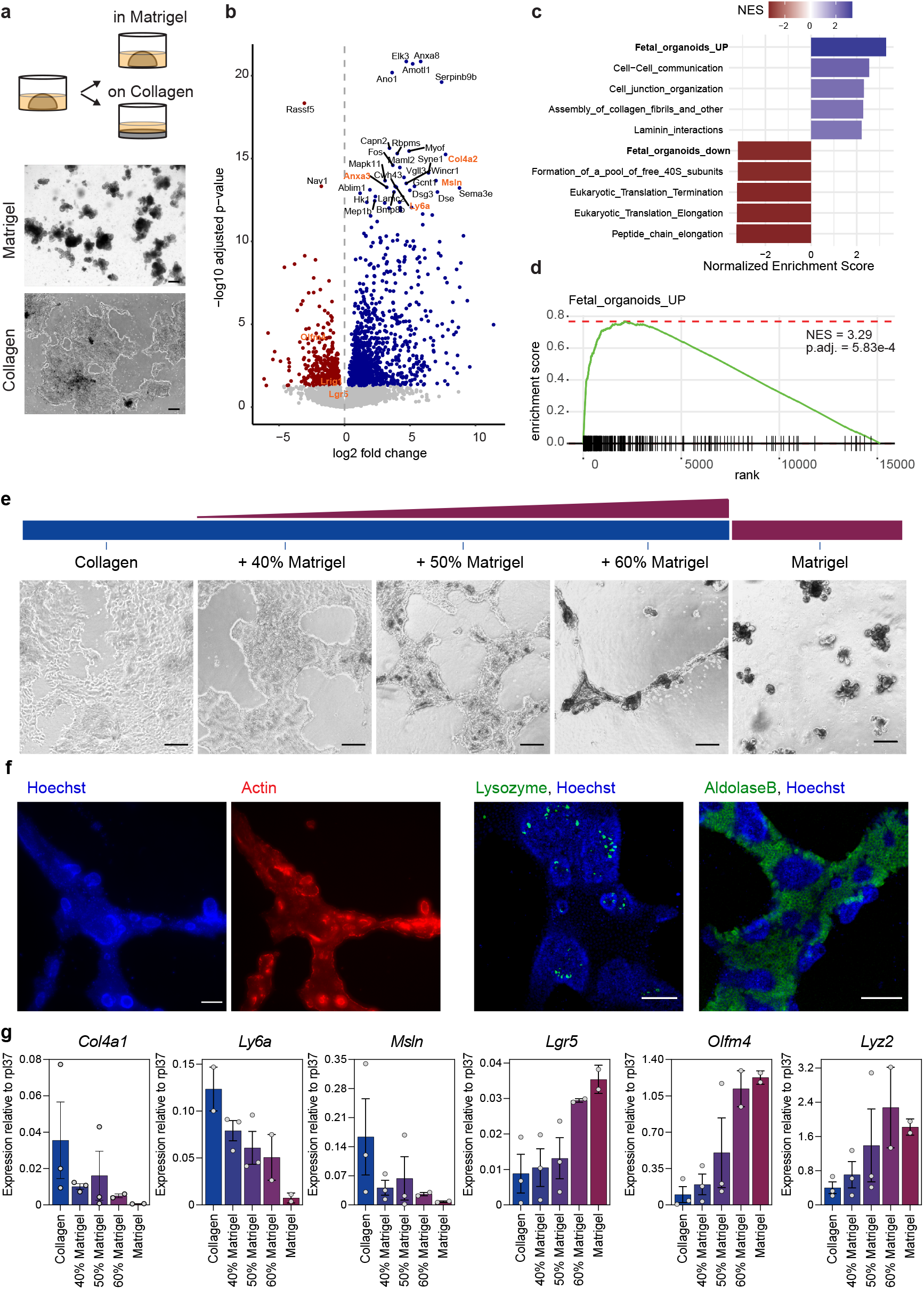
ECM controls intestinal stem cell genes and fetal-like genes. **a**, Intestinal cells from adult wild type mice were isolated and cultured for 2 passages in Matrigel. Matrigel derived organoids were then seeded on a collagen layer for at least 2 passages before RNAseq (n = organoids from 3 different mice), scale bar = 200um. **b**, Volcanoplot shows significantly (padj <0.05) upregulated (blue) and downregulated (red) genes in collagen vs. Matrigel cultures. The 30 most significant deregulated genes and intestinal stem cell genes (*Lgr5, Olfm4, Lrig1*) are labelled. Fetal-like genes and intestinal stem cell genes are highlighted in orange. **c**, Geneset enrichment analysis was performed with reactome pathway signatures and fetal-organoid signature (13). The top 5 up- and down-regulated pathways (ordered by NES) are shown. **d**, Fetal signature is highly enriched in collagen cultures. **e**, Collagen cultures were plated on different matrices, collagen with/without addition of Matrigel or pure Matrigel, scale bar = 100 um. **f**, Immunofluorescence of collagen+50% Matrigel shows crypt-like protrusions with apical accumulation of Actin, containing Lysozyme+ Paneth cells and were surrounded by AldolaseB+ enterocytes, scale bar = 100 um. **g**, qRT-PCR shows decrease of fetal-like genes (*Col4a1, Ly6a, Msln*) and upregulation of intestinal stem cell genes (*Lgr5, Olfm4*) and Paneth cell genes (*Lyz2*). Each dot represents an independent experiment, bar height = mean, error bar = s.e.m.

Recently synthetic matrices have been described that allow stem cell survival and proliferation or differentiation by changing the stiffness of the hydrogels (17), indicating that physical properties influence the epithelial cell composition. To avoid stiffness-related effects, we chose the collagen type I concentration that showed a similar stiffness as Matrigel **(Extended Data Fig. 1a)**. Similar to the Matrigel cultures, collagen cultures required R-spondin for long-term culture **(Extended Data Fig. 1b)**.

We performed RNAseq to compare the effect of the distinct ECM on the epithelial cells and detected marked differences in the gene expression profile **(Fig. 1b)**. Several of the most significant upregulated genes in the collagen cultures were characteristic of fetal organoids (13)(e.g. *Anxa3, Ly6a/Sca1, Msln, Col4a2*). In contrast, adult intestinal stem cell genes were downregulated (e.g. *Olfm4, Lgr5*). To analyse the differentially expressed genes in an unbiased manner, we performed gene set enrichment analysis. This confirmed a striking upregulation of fetal-organoid associated signatures in the collagen cultures, and a significant downregulation of adult organoid genes **(Fig. 1c, d)**, confirming earlier results in a 3D collagen system with the addition of Wnt3a (12).

Importantly, the effect of ECM components on gene expression is not due to difference in morphology (2D collagen vs. 3D Matrigel), but truly defined by the type of ECM as cells grown in a 3D collagen environment were transcriptionally similar to the 2D collagen cultures **(Extended Data Fig. 1c-e)**. Specifically, the fetal signature was highly upregulated in both collagen cultures (2D and 3D), and adult genes were downregulated. Interestingly, addition of Wnt3a, did not influence the expression of fetal genes, but downregulated several genes of the adult organoid signature, which were mostly associated with Paneth cells **(Extended Data Fig. 1d)**. Principal component analysis of the global transcriptome showed that the collagen I cultures cluster together, independent whether grown in 2D or 3D **(Extended Data Fig. 1e)**.

To further investigate the direct influence of ECM composition on the intestinal stem cells and expression of fetal-like genes we plated collagen-grown cells on an ECM with increasing concentrations of Matrigel **(Fig. 1e)**. Cells plated on collagen alone always grew as a flat 2D layer. We then mixed increasing concentrations of Matrigel into the collagen mixture, while maintaining the concentration of collagen I in the matrix. Strikingly, we observed crypt-like protrusions emerging from the epithelial network. These protrusions were enriched for Actin on the apical membrane and contained Lysozyme positive Paneth cells **(Fig. 1f)**, highlighting a crypt-like morphology and compartmentalization. Moreover, the flat epithelium surrounding the crypt-like structures accommodated Aldolase B positive enterocytes **(Fig. 1f)**. In line with the morphological changes, we observed that increasing concentrations of Matrigel reduced the expression of fetal-like genes and increased Paneth cell and stem cell gene expression in a dose-dependent manner **(Fig. 1g)**, despite the presence of the same collagen I concentration. Interestingly, we did not observe a similar change in absorptive enterocyte or goblet cell gene expression (*Alpi, Muc2*, respectively), nor an overall change in Wnt pathway activation (*Axin2*) when changing the ECM composition **(Extended Data Fig. 1f)**. This suggests that the increased basement membrane components have a dominant effect over the collagen resulting in induction of a crypt-like morphology and increased stem and Paneth cells.

### ECM components impact intestinal cell populations

To gain further insight into the different cell populations, we performed single cell RNAseq of collagen and Matrigel cultures. The cells present in Matrigel cultures could be clustered into nine distinct groups that could be related to known cell lineages of the small intestine. The heatmap shows the most differentially expressed genes in each cluster, and characteristic genes are highlighted that were used to assign the most probable cell type to each cluster. A large proportion of cells were characterised by expression of *Lgr5* as well as proliferation genes (Lgr5 stem cell and transit amplifying (TA-) cell clusters) **(Fig. 2a)**. Three clusters belong to absorptive enterocytes, probably marking different stages of development as has been previously observed *in vivo* (18) **(Extended Data Fig. 2a - *Apoa1*)**. Even rare enteroendocrine cells were identified as a distinct cell cluster in our data **(Extended Data Fig. 2a - *Chga*)**. The heatmap also highlights several clusters belonging to proliferative progenitors that are in the process of differentiation, e.g. early absorptive enterocytes and secretory cell progenitors **(Fig. 2a)**.

**Fig. 2.**
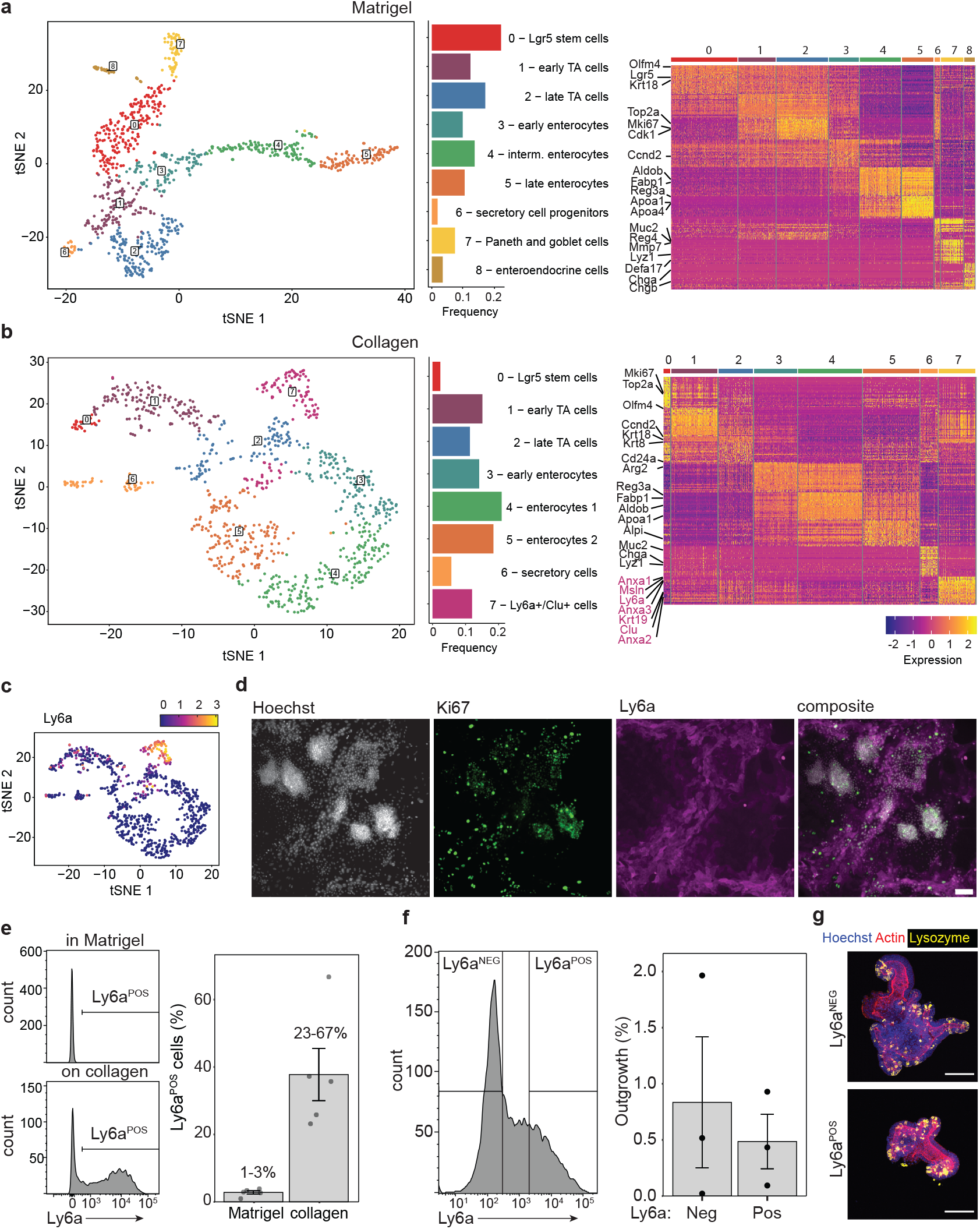
Distinct fetal-like cell population in collagen cultures with stem cell capacity. **a**, Clustering of Matrigel derived organoids at day 4 after passaging. Frequency plot shows proportion of cells assigned to each cluster. Heatmap shows top 30 differentially expressed genes in each cluster and labels of characteristic genes of intestinal cell types. **b**, Clustering of collagen derived cells at day 4 after passaging. Frequency plot shows proportion of cells assigned to each cluster. Heatmap shows top 30 differentially expressed genes in each cluster and labels of characteristic genes of intestinal cell types. Fetal-like genes are labelled purple. Cells for Matrigel and collagen scRNAseq were derived from the same organoid line. **c**, Ly6a expression in tSNE of collagen cultures. **d**, IF of collagen culture, stained for nuclei (Hoechst), proliferation (Ki67) and fetal-like gene (Ly6a). Scale bar = 100 um. **e**, Flow Cytometry of Ly6a-APC in Matrigel and collagen cultures. Each dot represents an independent experiment, bar height = mean, error bar = s.e.m., n = 5. **f**, Ly6a negative and positive cells from collagen cultures were sorted and plated in Matrigel. Organoids counted after 1 week, each dot represents an independent experiment, bar height = mean, error bar = s.e.m., n = 3. **g**, Immunofluorescence of organoids derived from Ly6a sorted cells, stained for nuclei (Hoechst - blue), Actin (red) and Paneth cells (Lysozyme - yellow). Scale bar = 100 um.

Intriguingly, in the collagen cultures, eight different cell clusters were distinguished which represented most of the cell lineages that are present in Matrigel **(Fig. 2b)**. However, we observed fewer stem cells, TA cells, and cells of secretory cell lineages **(Extended Data Fig. 2b)**. Due to their reduced number in this sample, enteroendocrine cells and Paneth/goblet cells form a single cluster but were reliably detected by expression of the characteristic genes in these cell types **(Extended Data Fig. 2b - *Chga* and *Lyz1/Muc2*)**. The distinct composition was further confirmed by immunofluorescence, which also revealed a distinct spatial distribution of Paneth cells and enterocytes in the collagen cultures **(Extended Data Fig. 3a)**.

Strikingly, one unique cluster was found on collagen that is not present in the Matrigel cultures. This cluster was characterised by fetal-like genes *Ly6a/Sca1, Clu, Msln, Anxa1* and *Anxa3* **(Fig. 2b**,**c and Extended Data Fig. 3b)**. Importantly, expression of these genes were restricted to this cluster, and were not detected in the other cell types **(Extended Data Fig. 3b)**. Interestingly, analysis of the combined cells from collagen and Matrigel cultures showed that all the secretory cells, e.g. Paneth and goblet cells, cluster together respective of their cell type, independent of whether they originate from collagen or Matrigel cultures. This suggests a similar transcriptional and differentiation profile in these cells, unaffected by the ECM composition **(Extended Data Fig. 3c)**. We next focused on the fetal-like cell cluster found only in the collagen cultures. Staining for Ly6a showed that these cells are located around the densely packed cell regions that contain Paneth and Lgr5+ stem cells, indicating a spatially distinct cell-type. Notably, Ly6a+ cells as well as the densely packed cell regions stained positive for the proliferation marker Ki67 **(Fig. 2d)**, suggesting that Ly6a- and Ly6a+ cells both sustain proliferation of the cultures. Analysis of the collagen and Matrigel cultures via flow cytometry confirmed that collagen cultures contained a reduced number of Lgr5-GFP cells and a large fraction of Ly6a+ cells **(Extended Data Fig. 3d)**. Overall we observed Ly6a+ cells in varying numbers in the collagen cultures, but consistently at negligible low percentages in the Matrigel cultures **(Fig. 2e)**. Furthermore, we did not observe any overlap of Lgr5-GFP and Ly6a staining, confirming that the Ly6a+ cells are a truly distinct cell population **(Extended Data Fig. 3e)**. Ly6a/Sca1+ as well as Clu+ cells have recently been described to be involved in the repair of the intestinal epithelium after damage and irradiation (12, 19). To test whether this *in vitro*-derived cell population has a similar stem cell potential, we sorted Ly6a-positive and -negative cells from collagen cultures and embedded them in Matrigel. Both cell populations had a similar capacity to form organoids **(Fig. 2f)**, could be passaged and were morphologically indistinguishable from each other. Importantly, the Ly6+ derived organoids also contained Paneth cells and showed that the Ly6+ cells have true stem cell capabilities **(Fig. 2g)**.

### Laminin controls intestinal stem cell fate

Having established that Matrigel and collagen affect the number of intestinal stem cells and fetal-like cells in organoid cultures, we asked which component of Matrigel is responsible for the observed effect. Matrigel is a basement-membrane extract and consists mainly of laminin and collagen IV. As the most striking morphological effect was observed when 50% Matrigel was added to the collagen hydrogel, we tried to reproduce this effect by providing either laminin or collagen IV separately **(Fig. 3a)**. Laminin was able to induce the same morphological as well transcriptional response as Matrigel in a dose-dependent manner **(Fig. 3b)**. In contrast, addition of collagen IV, did not result in any morphological or transcriptomic changes **(Fig. 3c)**. These data indicate that laminin is the main component responsible for the induction of the crypt-like structures and expression of genes associated with adult intestinal stem cells.

**Fig. 3.**
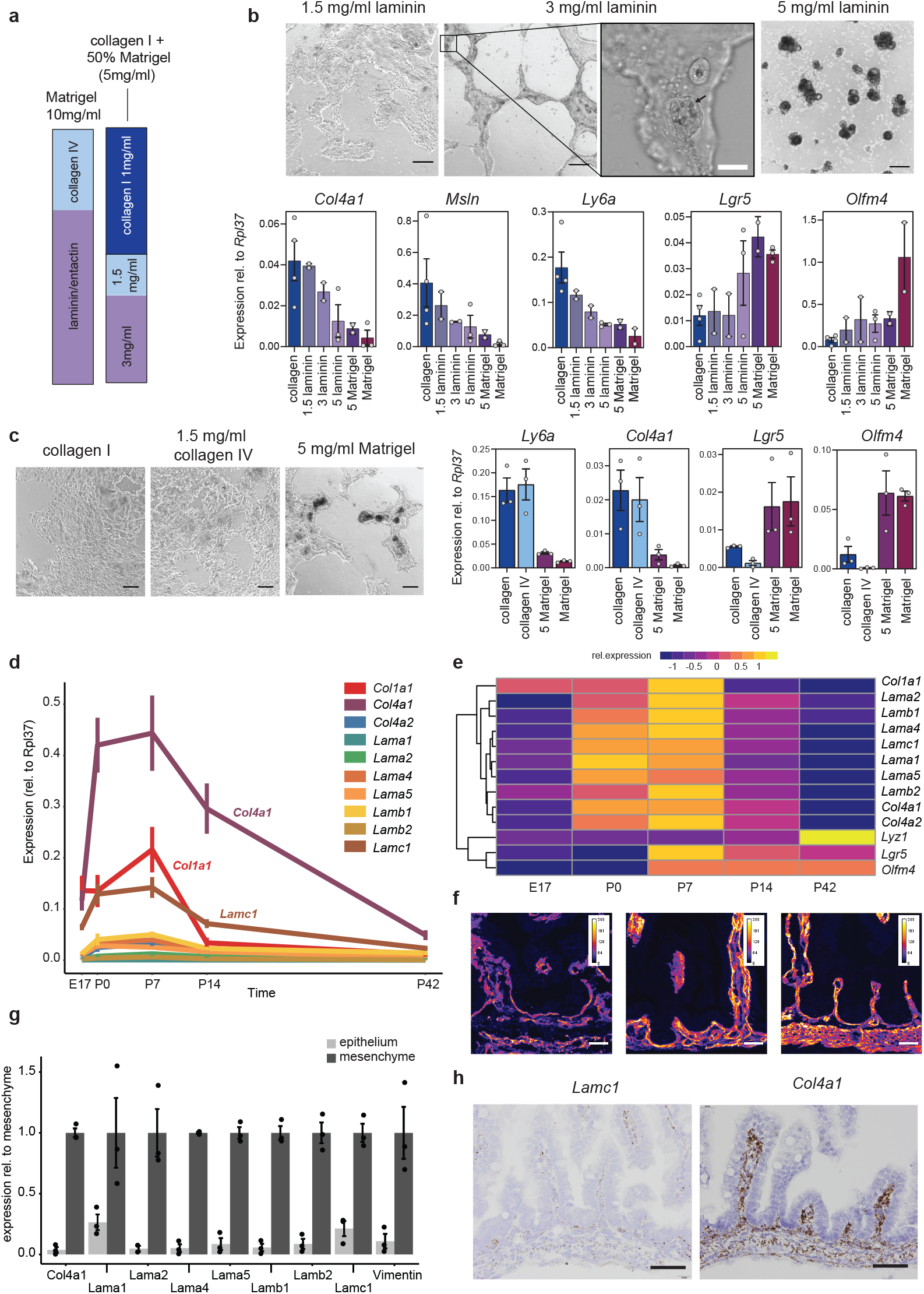
Laminin induces crypt-like structures and is upregulated during crypt formation in vivo. **a**, Matrigel mainly consists of Laminin and Collagen IV. Single basement membrane components (laminin/entactin or collagen IV) were added to a 1mg/ml collagen hydrogel. **b**, Cells grown on collagen were plated on collagen I + laminin (1.5 mg/ml, 3 mg/ml, 5mg/ml) or + Matrigel (5 mg/ml), or collagen or Matrigel alone. Laminin induces morphological changes, as well as a similar downregulation of fetal-like genes (*Col4a1, Ly6a, Msln*) and upregulation of stem cell markers (*Lgr5, Olfm4*), n = 2-4 independent samples per mixture, black scale bar = 200um, white scale bar = 50um. Black arrow indicates Paneth cells in crypt-like protrusion. **c**, Cells grown on collagen were plated on collagen I, collagen I + 1.5 mg/ml collagen IV, collagen I + 5 mg/ml Matrigel or Matrigel alone. n = 3. Scale bar = 100um. **d**, qRT-PCR analysis of intestinal tissue at different timepoints during development, N=5 mice per time point, height = mean, error bar = s.e.m. **e**, Heatmap of qRT-PCR averages scaled for each gene showing that all basement membrane genes cluster together, separate from collagen I (*Col1a1*) and are specifically upregulated at P0 (birth) and P7. Intestinal stem cell genes (*Lgr5, Olfm4*) are upregulated at P7. Paneth cell genes (*Lyz1*) appear later at P14, and increase further until P42. **f**, Laminin immunofluorescence shows increased laminin levels during crypt development, n = 3 mice, scale bar = 45um. **g**, Intestinal crypts (without villi) at P7 was EDTA-separated for epithelium and mesenchyme. *Vimentin* shows enrichment of mesenchymal cells that are absent in the epithelial fraction. Laminin and collagen IV genes are highly expressed in the mesenchyme with little contribution of epithelial cells, Error bars = s.e.m. **h**, RNA in situ hybridization (RNAscope) for Lamc1 and Col4a1 at P0 shows predominant mesenchymal expression, scale bar = 50um. For all bar graphs (b, c, g), each dot represents an independent experiment, bar height = mean, error bar = s.e.m.

We noticed parallels between the developing small intestine *in vivo* and our *in vitro* system. Before birth, the mouse small intestine only contains villi and proliferating intervillus structures. Cells from both of these compartments can contribute to the pool of adult stem cells after birth (20). Around 2 weeks after birth, the final crypt structures have formed and mature Paneth cells can be detected (21– 23). Therefore we wondered if the development of intestinal crypts *in vivo* showed a similar change in the ECM. We analysed the small intestine of wild type mice from E17 until 6 weeks after birth (P42). Intriguingly, a dramatic change around birth in the expression of all laminin isoforms (*Lama, Lamb* and *Lamc* genes) as well as collagen IV (*Col4a1, Col4a2*) was observed **(Fig. 3d)**. This spike in expression could be detected for one week, after which the levels of expression reduced. The expression of collagen type I (*Col1a1*) only slightly changed and did not follow the same expression burst as observed with the basement membrane components **(Fig. 3d, e)**. Immunofluorescence staining for Laminin further validated an increase in protein levels from E19 to P7 **(Fig. 3f)**. Next, we aimed to determine which cells are responsible for the increased expression of the basement membrane components. We separated the epithelium from the mesenchyme of wild type mice at one week after birth (P7). All of the basement membrane components were predominantly expressed in the mesenchyme and only little expression was found in the epithelium **(Fig. 3g)**. RNA *in situ* hybridization for laminin γ1 (*Lamc1*) and collagen IV (*Col4a1*) confirmed a specific upregulation exclusively in the mesenchyme **(Fig. 3h and Extended Data Fig. 4a)**. To visualise the laminin network, we cleared and stained the small intestine of a mouse one week after birth (P7). Laminin was detected surrounding the nascent crypts as well as villi **(Extended Data Fig. 4b)**. This indicates that the dramatic remodelling of the ECM during crypt formation is driven by the mesenchyme and not the epithelium.

### Laminin signalling controls intestinal epithelium change via ITGA6

Our data revealed a potent induction of the adult stem cell program and morphological changes by the addition of laminin and that formation of intestinal crypts in gut development coincides with a temporal increase in laminin production by the mesenchyme. To gain further insight into the molecular signalling between the epithelium and the basement membrane components, we evaluated the expression of their receptors during development. The key family of receptors for the ECM are integrins and dystroglycan receptors. We analysed a publicly available dataset of intestinal epithelial cells during development (24) for expression changes in integrins and dystroglycan encoding genes. Strikingly, at embryonic day E18, a specific upregulation of *Itga6, Itgb4, Dag1* and *Itga3* was observed **(Fig. 4a)**, whereas all other integrins were only lowly expressed and did not change during development **(Extended Data Fig. 4c)**. ITGA6 (together with ITGB4) is a known receptor for Laminin (25). RNA *in situ* hybridization for *Itga6* confirmed the previous analysis and showed a marked upregulation of *Itga6* after birth **(Fig. 4b)**. Remarkably, the expression is even more pronounced in the intervillus region and stays enriched in the crypts compared to villi. Next we asked whether ITGA6 was mechanistically involved in laminin-induced phenotypic change. Blocking ITGA6, in our previously described *in vitro* assay, diminished the induced morphological changes, prevented the upregulation of intestinal stem cell genes and impaired the downregulation of fetal-like genes **(Fig. 4c)**. Given this central role of laminin signalling during development, we evaluated the importance of laminin-epithelial signalling in the adult intestine *in vivo*. The laminin γ-chain is common to all laminin heterotrimeric proteins and was the highest expressed laminin gene during development **(Fig. 3d)**. We crossed the *Lamc1* flox mouse to a ubiquitously expressing cre mouse, to delete laminin expression in adult mice as previously reported **(Fig. 4d)** (26). Based on our *in vitro* model, we predicted that depletion of laminin would result in a decrease of stem cells and Paneth cells, as well as an upregulation of fetal-like genes. Indeed, 3 weeks after deletion of *Lamc1*, the intestinal epithelium showed a marked downregulation of *Lgr5, Olfm4* and *Lyz1* expression and upregulation of fetal-like genes in line with our *in vitro* experiments **(Fig. 4e, f)**.

Overall these data support the hypothesis that the ECM composition, as orchestrated by the mesenchyme, induces and maintains crypt formation as well as intestinal stem cell specification in the developing and adult intestine **(Fig. 5)**.

**Fig. 4.**
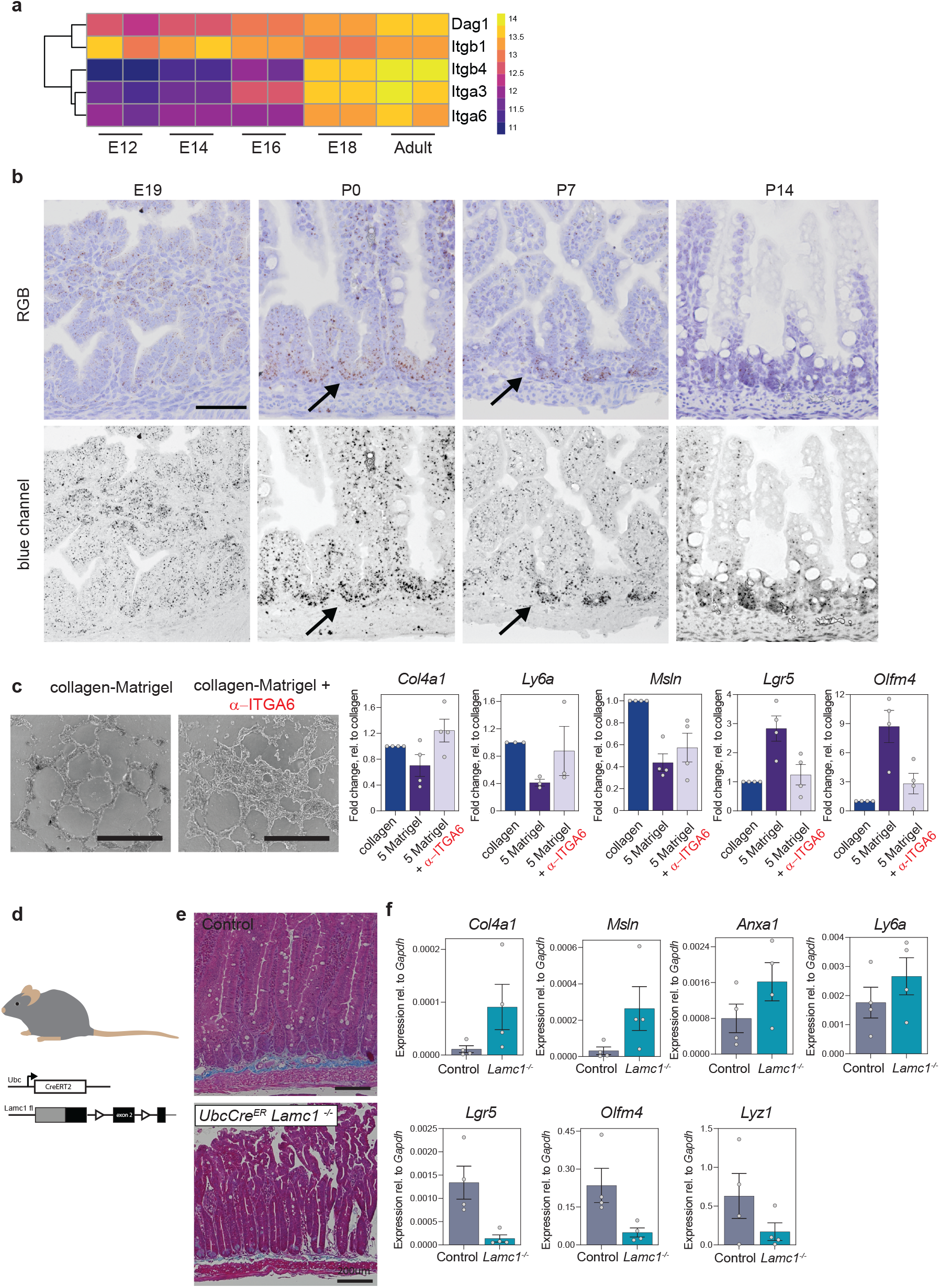
Laminin signalling, via Integrin alpha 6, induces crypt formation in vitro and controls intestinal stem cells in vivo. **a**, An RNAseq dataset (GSE115541 (24)) consisting of epithelial intestinal cells during development shows specific upregulation of laminin receptor encoding genes *Itga6, Itga3, Itgb4, Dag1* from E18 onwards. **b**, RNA *in situ* hybridization for *Itga6* shows increased expression at P0, with accumulation at the intervillus region. Bottom row shows blue channel images for increased contrast of the RNA *in situ* signal. Black arrows indicate inter-villus region and nascent crypts. Scale bar = 50um. **c**, Cells were grown on collagen and then seeded on collagen I or collagen I + 5 mg/ml Matrigel (5 Matrigel) mixture with or without addition of ITGA6 neutralising antibody (*α*-ITGA6). Inhibition of ITGA6 blocks downregulation of fetal-like markers and blocks increase of stem cell markers. Each dot represents an independent experiment, n = 4. All values are relative (fold change) to its collagen control, scale bar = 1000 um. **d**, *Ubc-CreERT2 Lamc1loxP/loxP* adult mice (>6 weeks old) were injected with tamoxifen to delete *Lamc1* throughout the body, and mice were analysed 21 days after injection. **e**, Trichrome staining shows epithelial phenotype after deletion of *Lamc1*, scale bar = 200um. **f**, Fetal-like genes were upregulated whereas stem cell and Paneth cell markers were downregulated after deletion of *Lamc1*. Each dot represents independent experiment, n = 4 mice per group, bar height = mean, error bar = s.e.m.

**Fig. 5.**
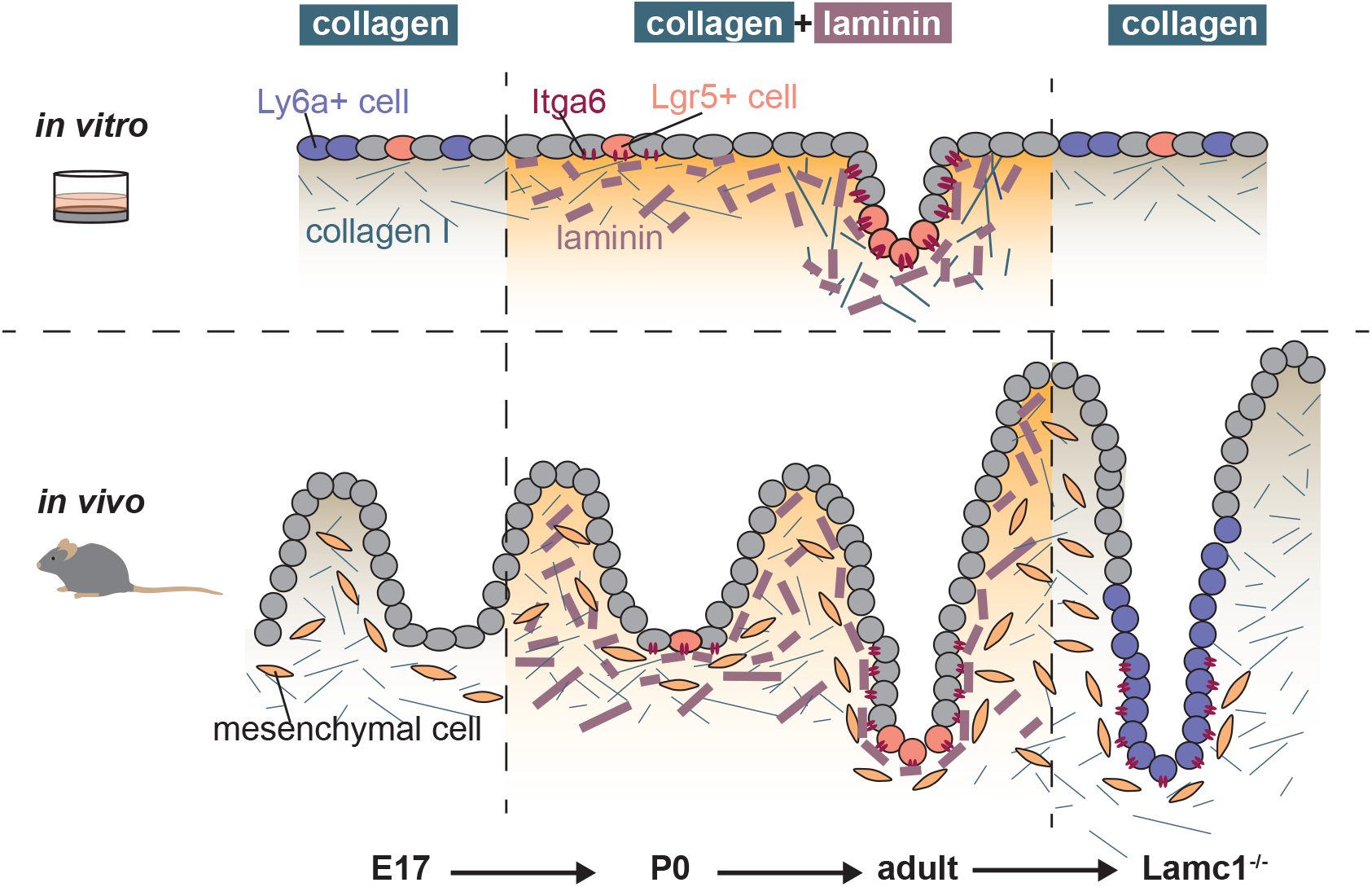
Graphical abstract. Parallels between *in vitro* cultures with different ECMs and *in vivo* crypt formation and homeostasis. *In vitro*, adult intestinal cells contain fetal-like cells (Ly6a+) and few intestinal stem cells (Lgr5+) and grow exclusively in a 2D layer. Addition of laminin induces an increase of Lgr5+ cells and Paneth cells (via ITGA6), resulting in a morphological crypt formation. *In vivo*, the mesenchyme dramatically remodels the ECM and increases laminin concentration. Concurringly, epithelial cells in the inter-villus region upregulate expression of the laminin receptor ITGA6 and progress to form intestinal crypts. Even in the adult, intestinal crypts rely on the laminin signalling, as depletion of laminin results in a fetal-like epithelium.

## Discussion

In recent years the focus on the intestinal niche shifted considerably. With the development of the organoid technology, the main focus has been on the Paneth cells as the key part in establishing an intestinal stem cell niche (2, 27). Careful analysis *in vivo* however showed that Paneth cells are dispensable, and indicated that mesenchymal cells were the main provider of the intestinal niche (28–30). Furthermore, even organoid cultures relied on the addition of growth factors that are usually secreted by the mesenchyme (e.g. Rspondin). More recently, the focus shifted onto this mesenchymal niche and the different subtypes of fibroblasts that constitute this niche *in vivo* (1). Next to secreted growth factors, such as Wnt ligands and R-spondin, data support a strong role for these mesenchymal cells in controlling the composition of the extracellular matrix *in vivo* and therefore points to a potential role for for the creation of a niche (8, 9, 31, 32). Here we show that both, the interstitial matrix (collagen I) as well as the basement membrane matrix (Matrigel/laminin) support the long term growth of the intestinal epithelium with the presence of all major cell lineages. However, morphological crypt like structures were only observed when the ECM contained high concentration of laminin (and not collagen IV), supporting previous findings in other *in vitro* hydrogel settings (33). This is despite the presence of Lgr5+ stem cells and Paneth cells in the collagen cultures, which are shown to be responsible for crypt-budding in Matrigel grown organoids (34). An interesting question that arises, whether collagen induces the fetal-like reprogramming or the lack of laminin signalling. In a recent study from the Liberali lab the authors showed that organoid growth in Matrigel from single cells undergoes a similar, transient fetal-like cell state which then declines as Lgr5+ intestinal stem cells and Paneth cells appear (34). This could indicate that cells taken out of their ECM environment need some time to adapt to the laminin-rich environment and hence undergo a temporarily fetal-like state. Therefore a transient disruption of laminin-signalling might already induce the fetal-like response. Also our data indicate, increasing concentations of laminin revert the fetal-like state in the collagen cultures, despite an unchanged collagen concentration.

Overall our data support the hypothesis that the basement-membrane produced by the epithelium is not sufficient to increase stem cell numbers and induce a morphological crypt formation, and therefore extra-epithelial cells (e.g. fibroblasts) need to remodel the ECM prior to crypt formation. It seems likely that pericryptal fibroblasts are responsible for supplying the epithelium with laminin due to their close proximity to the crypt epithelium. These cells (Foxl1+, Gli1+, Pdgfra+ and/or CD34+ fibroblasts) also secrete Wnt ligands and R-spondins, which are essential for crypt homeostasis and would therefore express all necessary factors for the intestinal niche (35–39).

We saw a dramatic increase in expression of basement membrane components in the mesenchyme around birth, supporting earlier observations (8, 9). This coincides with the expression of *Itga6*, a known laminin receptor, in the intervillus cells and nascent crypts (25, 32). We detected also upregulation of other integrins (e.g. *Itga3*) *in vivo* and we would speculate that deletion of *Itga6* over longer time periods, would be compensated by other integrins. A knock-out mouse of *Itga6* has been reported and develops normal crypts, however is susceptible to colitis and adenoma formation at later stages (40). Nevertheless, we were able to interfere with the ITGA6 receptor *in vitro* by use of a neutralising antibody, highlighting its central role.

In recent years, much effort came to define the mechanical properties for stem cell proliferation and cell compartmentalization (17, 41). Our study showed that a changed ECM with a similar soft stiffness, affects the epithelium via laminin signalling through the ITGA6 receptor and points away from a pure physical explanation of the matrix (e.g. changed stiffness) as the responsible factor for the observed effects. Therefore a localized ECM-epithelial cell signalling is potentially more crucial for stem cell specification and crypt formation. Overall, our study highlights a central role of the ECM in controlling intestinal stem cells and crypt morphology during development and in the adult.

## Materials and Methods

### Mice and crypt isolation

Crypts were isolated from wild type and Lgr5–EGFP–ires–CreERT2 mice (42) following previously described methods (2). In vivo experiments performed in Amsterdam were approved by the Animal Experimentation Committee at the Academic Medical Center in Amsterdam (ALC102556 and Lex 274, WP 17-1884-1-01) and performed according to national guidelines. For Lamc1loxP mouse experiments, husbandry and research protocols were carried out in accordance with protocol AUA00003140, which was approved by The Medical College of Wisconsin’s Institutional Animal Care and Use Committee (IACAUC). Ubc-CreERT2 (43, 44) Lamc1loxP/loxPl (45)mice were induced with four intraperitoneal injections of 1 mg/ml tamoxifen on four consecutive days. Tissue of mice was harvested 21 days after first injection.

### Organoid growth in Matrigel

Small intestinal crypts were isolated and cultured as previously described (2). Isolated crypts (via EDTA 2mM) were embedded in growth-factor reduced Matrigel (Corning 356231). Medium (ENR) was supplemented with EGF [50ng/ml], conditioned media of Noggin (10%) and conditioned media of Rspo1 (20%). For the first 48hrs after initial purification, media was supplemented in addition with CHIR-99021 [5uM] and Rock inhibitor, Y-27632 [10uM]. Culture medium was changed every other day. Wn3a-conditioned media was produced from L-Wnt3a cells in 10% FBS and was added at 50% to ENR medium (WENR).

### Cell growth on collagen

Mouse small intestinal cells were grown on collagen as previously described (10). First, mouse intestinal organoids grown in Matrigel were released from Matrigel by incubation with Cell Recovery solution (BD Science #354253) for 30 mins on ice and then mechanically dissociated before being placed on a collagen layer. Collagen Rat tail type I (Corning 354236) layer was prepared and neutralized to a concentration of 1mg/ml. The collagen mixture was added to a 6 well plate (1ml/well) and left to gellate at 37C for 1h. Cells collected from matrigel cultures were resuspended in medium and plated on top of the hydrogel. For the initial 48hrs after passaging, the medium was supplemented with Rock inhibitor, Y-27632 [10uM]. Medium was changed every other day. Cells were collected 6 days after plating in 1 ml PBS and incubated with 60-80 ul collagenase IV [10mg/ml] (Sigma #C5138-1G) at 37 C for 10 mins, then washed 3 times in 10 ml PBS. Cell pellets were resuspended in culture medium and split into a ratio of 1 well/2-3 wells.

### Cell growth on collagen: basement membrane components

Higher concentrations of Collagen IV (Corning #354233, 1 mg/ml) were achieved by freeze-drying the sample in a lyophilizer (ZIRBUS technology) at 0.05 mbar and -80C for a period of 72h. The dried collagen IV was dissolved in Phosphate Buffered Saline solution at a final con-centration of 2 mg/ml. Collagen I and basement membrane components, Matrigel, laminin (Thermo Fisher #10256312) or Collagen IV, were mixed to create desired concentrations and added to a 12 well plate (0.5 ml/well), final concentration of collagen I was 1mg/ml. Cells grown on collagen I (passage number 2-5) were collected and washed following the previously mentioned procedure. Pellet was resuspended in 4 ml of medium and 1 ml was plated on top of each well. For the initial 48h after passaging, medium was supplemented with Rock inhibitor, Y-27632 [10uM]. Medium was changed every other day (ENR). Cells were collected 4 days after passaging in 1 ml PBS and incubated with 60 ul collagenase IV [10mg/ml] for 10 mins at 37 C, then placed on ice for 40 mins before pelleting. For blocking of ITGA-6, collagen-grown cells were collected and incubated with ITGA-6 neutralizing antibody (Merck Millipore #MAB1378, clone NKI-GoH3 at 40ug/ml) for 60 mins at 37 C before plating on hydrogels. Medium was renewed every 48hrs (ENR) with newly added antibodies and RNA was harvested on day 4.

### Immunofluorescence

Immunofluorescence staining was used to reveal the cellular composition of Matrigel, collagen I and collagen:basement membrane cultures. Samples were fixed in 4% formaldehyde for 1h, permeabilized with 0.2% Triton-X in PBS for 1h and blocked with Antibody Diluent (ImmunoLogic #VWRKUD09-999) for 30 mins. The following antibodies were used, Lysozyme EC 3.2.1.17 (1:200, DAKO #A0099), AldolaseB EPR3138Y(1/200, abcam #ab75751), Ly6a-APC (1/200, eBioscience 17-5981-81). Snap frozen mouse intestinal tissue (E19, P0 and adult) were cut at a thickness of 10 um using a cryostat and transferred onto Superfrost microscope slides. Sections were fixed with 4% PFA for 10 mins at RT, permeabilized with 0.2% Triton-X100 for 10 mins and blocked with Antibody Diluent for 10 mins. Slides were incubated with Laminin Polyclonal Antibody (1/200, Thermo Fisher #PA5-22901) overnight at 4C. Images were taken using the Leica TCS SP8 X microscope.

### RNAscope

Formalin-fixed paraffin embedded (FFPE-) slides (5um) were prepared and stained according to the manufacturer’s instruction with RNAscope Detection kit 2.5 (brown) with the following probes, Mm-Lamc1 (ACD #517451) and Mm-Col4a1 (ACD #412871) and Mm-Itga6 (ACD #441701).

### Flow Cytometry and FACS

Cells grown on collagen were collected in 0.5ml PBS and incubated with 80ul of collagenase IV [10 mg/ml] for 10 mins at 37C. After 2 wash steps with PBS, the cells were dissociated into single cells by incubation with 1ml TrypLE Express for 15 mins at 37C with mechanical dissociation after 10 mins. Cells were then incubated with anti-mouse Ly6a-APC (1/1000, eBioscience #17-5981-81) for 30 mins on ice. For sorting and plating of cells, 10k cells/25ul Matrigel/well were plated and incubated with GSK-3 inhibitor, Chir-99021 [5um] and Rock inhibitor, Y-27632 [10uM] for the first 48hrs.

### Single cell RNAseq

Organoids grown in Matrigel and cells grown on collagen, both derived from the same organoid line, were collected 4 days after passaging and digested with TrypLE (Invitrogen) for 20 min at 37°C. Cells were passed through a cell strainer with a pore size of 20 µm. Cells were sorted (3000 single, living cells) by fluorescence-activated cell sorting (FACS/ SONY sorter), manually counted and adjusted, and processed using 10x Chromium Single Cell 3’ Reagent Kits v3 library kit. Reads were de-multiplexed, aligned to the GRCm38/mm10 reference genome (‘refdata-cellranger-mm10-3.0.0’) and counted using the 10x Genomics Cellranger software (v3.1.0). Raw count matrices were analysed with the Seurat library (46)in R. Cells expressing less than 1000 genes or more than 15% mitochondrial genes were removed.

### RNAseq

Organoids grown in Matrigel were derived from 3 individual mice (n = 3) and subsequently grown on collagen supplement with ENR or WENR, in collagen with WENR and in Matrigel supplemented with ENR or WENR and were collected for RNA isolation 4 days after passaging. The quality was assessed using the Agilent RNA ScreenTape Assay. RNA-seq libraries were prepared with the KAPA mRNA Hyper Prep Kit (KAPA Biosystems) using 550 ng of total RNA and sequenced on Illumina Hiseq4000 (single read, 50bp). The reads were aligned to the GRCm38/mm10 reference genome via Rsubread (47) and the reads quantified by featurecounts (48). The raw counts were further analysed with DESeq2 (49) and gene-set enrichment analysis was used with the fgsea package (50). RNAseq dataset of epithelial, fetal intestine, GSE115541 (24) was similarly analysed with DEseq2.

### Code and Data availability

All code and necessary processed data used for the analysis can be found at https://github.com/davidhuels/collagen_matrigel_project. The raw data from RNAseq can be found on Array Express E-MTAB-10082.

### Stiffness Young’s Modulus

Young’s modulus was determined with a displacement-controlled nanoindenter (Piuma, Optics 11). A spherical probe with a radius of 50 um and a cantilever stiffness of 0.5 N/m. All measurements were performed with a 1mm layer of hydrogel covered with PBS according to the manufacturer protocol.

### Tissue clearing (Ce3D)

Tissue clearing was realised following the Clearing enhanced 3D microscopy (Ce3D) protocol (51).Briefly, 20 ml of Ce3D clearing solution was prepared in a fume hood by dissolving Histodenz (86% (w/v), Sigma) in 40% N-methylacetamide (Sigma-Aldrich #M26305) in PBS. The mixture was placed at 37 C for 72h or until fully dissolved. Triton-X100 (0.1% v/v) and 1-thioglycerol (0.5% v/v, company) were added to the dissolved mixture. Mouse intestinal tissue (E19, P0 and adult) was isolated, cut open longitudinally, pinned and fixed overnight in 4% PFA at 4 C. Fixed tissues were incubated in a blocking buffer containing 1% bovine serum albumin and 0.3% Triton-X100 overnight on a shaker. Tissues were then incubated with Laminin (1:100, ThermoFisher #PA5-22901) and EpCAM (1:100, Biolegend #324221) in Ce3D solution for 72h on a shaker. Tissues were washed with PBS and incubated with Ce3D clearing solution for 48h on a shaker. Cleared tissues were incubated with Hoechst (1:1000) for 8h before mounting with Ce3D solution and imaging using the Leica TCS SP8 X microscope.

### qRT-PCR

Cells cultured in Matrigel, on collagen or on collagen:basement membrane hydrogels were collected 4 days after passaging following the previously described protocols. Mouse intestine whole tissue, mesenchymal fraction and epithelial fraction were isolated and digested following previously reported procedures. RNA was isolated using the NucleoSpin RNA kit (BIOKE #MN740955250) following the manufacturers’ guidelines. Reverse transcription was performed using the SuperScriptTM III First-Strand Synthesis Mix (Thermo Fisher #18080085)) on isolated RNA to obtain complementary DNA (cDNA). All quantitative reverse transcription-polymerase chain reactions (qPCR) were performed using the SYBR green detection system, each sample was run with 2 technical replicates. The primer sequences are provided in Supplemental Table 1.

### qRT-PCR of Lamc1 mouse model

Intestinal epithelium and mesenchyme were separated by incubation of tissue fragments in ice cold BSS buffer (1mM KCl, 96mM NaCl, 27mM Sodium Citrate, 8mM KH2PO4, 5.6mM Na2HPO3, 15mM EDTA) containing protease and phosphatase inhibitors at 4°C with vigorous shaking for 30minutes. Sheets of mesenchyme are then removed with forceps. Total RNA from epithelial cells was DNase treated with EZ-DNase (Invitrogen). Superscript VILO master mix (Invitrogen) was used to synthesize cDNA. TaqMan assays and TaqMan Gene Expression Master Mix (Applied Biosystems) were used to measure gene expression: LamC1 (Mm00711820_m1), Anxa1 (Mm00440225_m1), Col4a1 (Mm01210125_m1), Ly6a (Mm00726565_s1), Msln (Mm00450770_m1), Lyz1 (Mm00657323_m1), Olfm4 (Mm01320260_m1), and LGR5 (Mm00438890_m1). GAPDH (4351309) was used for normalization.

## ACKNOWLEDGEMENTS

DJH was funded by **EMBO ALTF 1102-2017**. JL was funded by the MRC MR/N021800/1 and OJS was funded by CRUK A17196 and A21139. Felipe A Vieira Braga helped with the preparation of cells for the scRNAseq. Kristin Komnick supported the work with the intestinal Lamc1 loxP tissues. We thank the technicians and consultants of the Core Facility Genomics of the Amsterdam UMC for performing and advising on the RNAseq and single cell RNAseq. We are thankful to Thijs Beldman and the Medical Biochemistry department of Amsterdam UMC for helping with the freeze-dryer and Fernanda M. Bosada-Musselwhite for the provision of embryos. The preprint version of this manuscript was made using the Henriques lab bioRxiv template.

## AUTHOR CONTRIBUTIONS

DJH and RR designed the project. RR, SvN, VMW, TMG, VM, JL, OJS, LV, JPM and DJH contributed to data acquisition, analysis and/or interpretation. DO, MB and KSC performed the experiments with the Lamc1 mouse. RR, JPM, LV and DJH wrote the manuscript. LV, JPM and DJH supervised the project.

## Supplementary Data

**Table S1.**
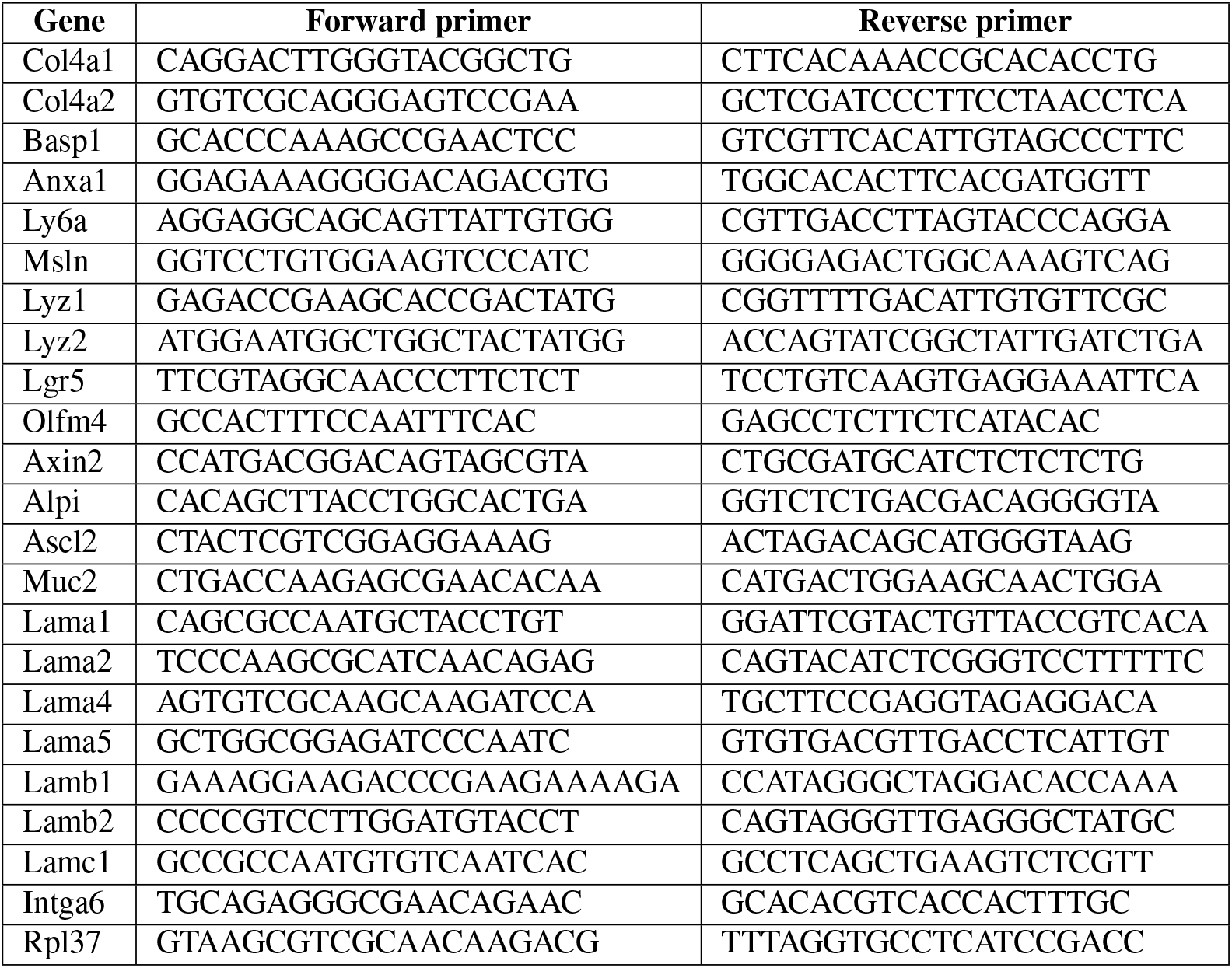
qRT-PCR primer list.

**Fig. S1.**
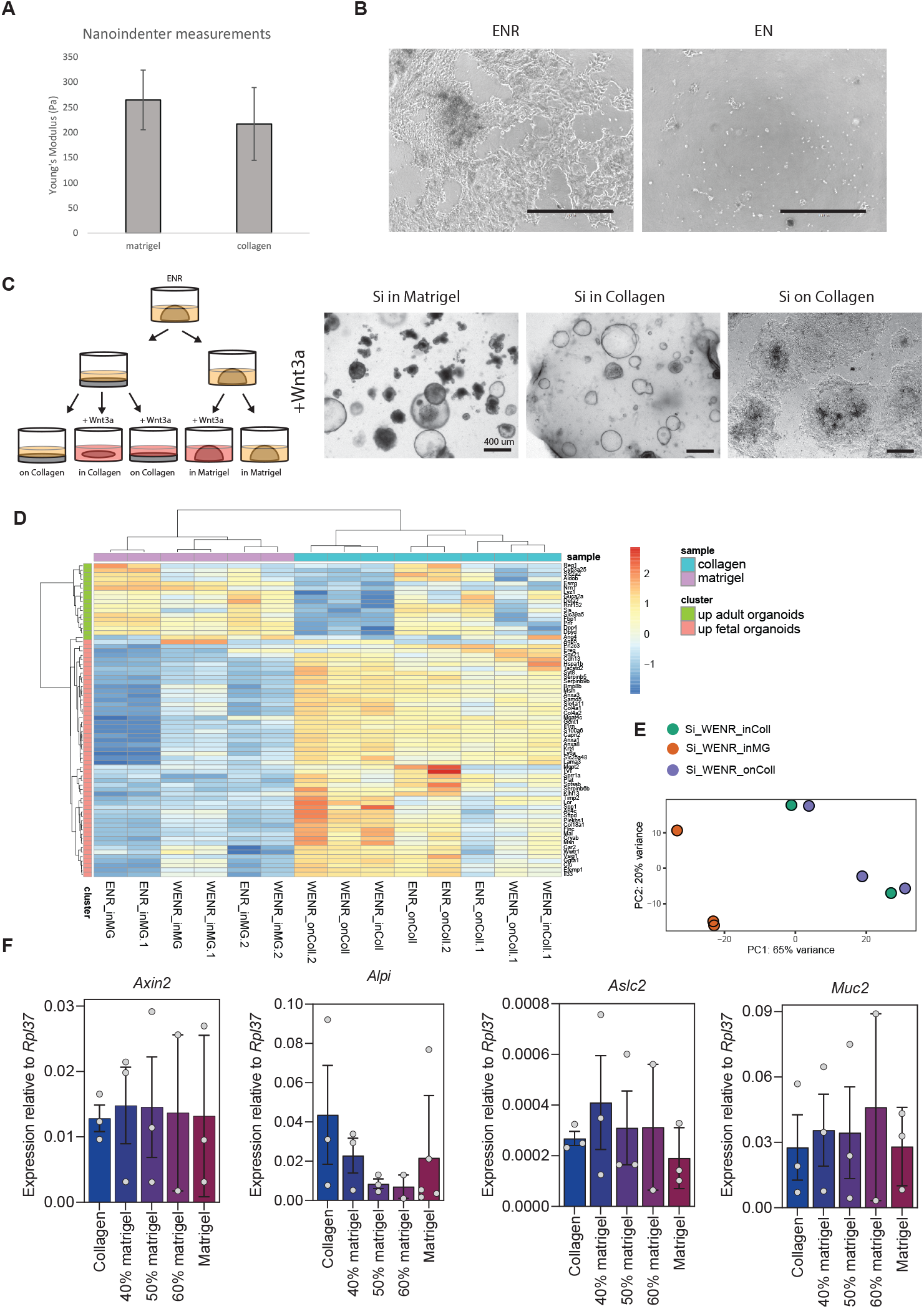
**a**, Stiffness measurement (Young’s modulus) was measured with a Nanoindenter (Piuma - Optics 11) on collagen or Matrigel hydrogel in PBS at room temperature. Technical measurements from the same sample were averaged, and repeated with new hydrogels (n=2). **b**, collagen cultures require the addition of R-spondin for long-term growth. Images after 1 passage with EN (EGF, Noggin) or ENR (EN+R-spondin). Scale bar = 1000um. **c**, Schematic of experimental design of RNAseq of 2D and 3D collagen cultures, with/without addition of Wnt3a-CM (50%). Scale bar = 400um. **d**, Heatmap of Mustata fetal gene signature (13), selected for highly variable genes (sd >1.5), shows that all collagen cultures are enriched for fetal-like genes, whereas Matrigel cultures are enriched for adult-genes. Addition of Wnt3a decreases expression of adult-genes in collagen cultures. ENR = EGF, Noggin, R-spondin. WENR = Wnt3a-CM (50%) + ENR. **e**, PCA of Matrigel culture, 2D and 3D collagen culture, all with addition of Wnt3a shows that collagen cultures cluster together, separated from the Matrigel culture, n = 2-3 samples per group, derived from different mice. **f**, Expression of Wnt target gene (Axin2), enterocyte gene (Alpi), stem cell marker (Ascl2) and goblet cell marker (Muc2). Each dot represents an independent sample, bar height = mean, error bar = s.e.m.

**Fig. S2.**
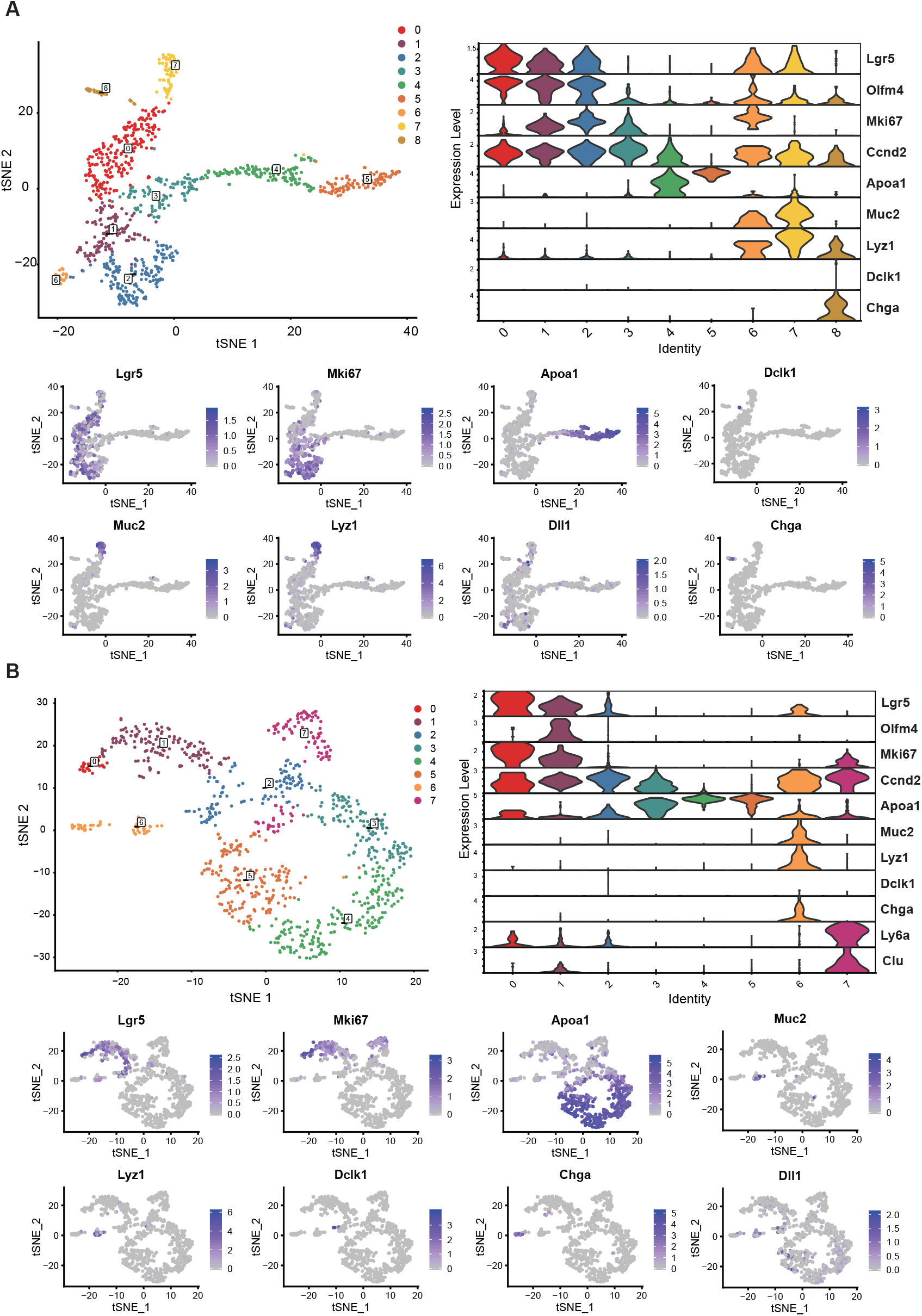
**a**, scRNAseq of Matrigel cultures. Clusters as described and as shown in Fig.2a. Violinplot shows expression of cell lineage genes per cluster. Expression of individual cell lineage genes overlaid in tSNE-plot. **b**, scRNAseq of collagen cultures. Clusters as described and as shown in Fig. 2b. Violinplot shows expression of cell lineage genes per cluster. Expression of individual cell lineage genes overlaid in tSNE-plot.

**Fig. S3.**
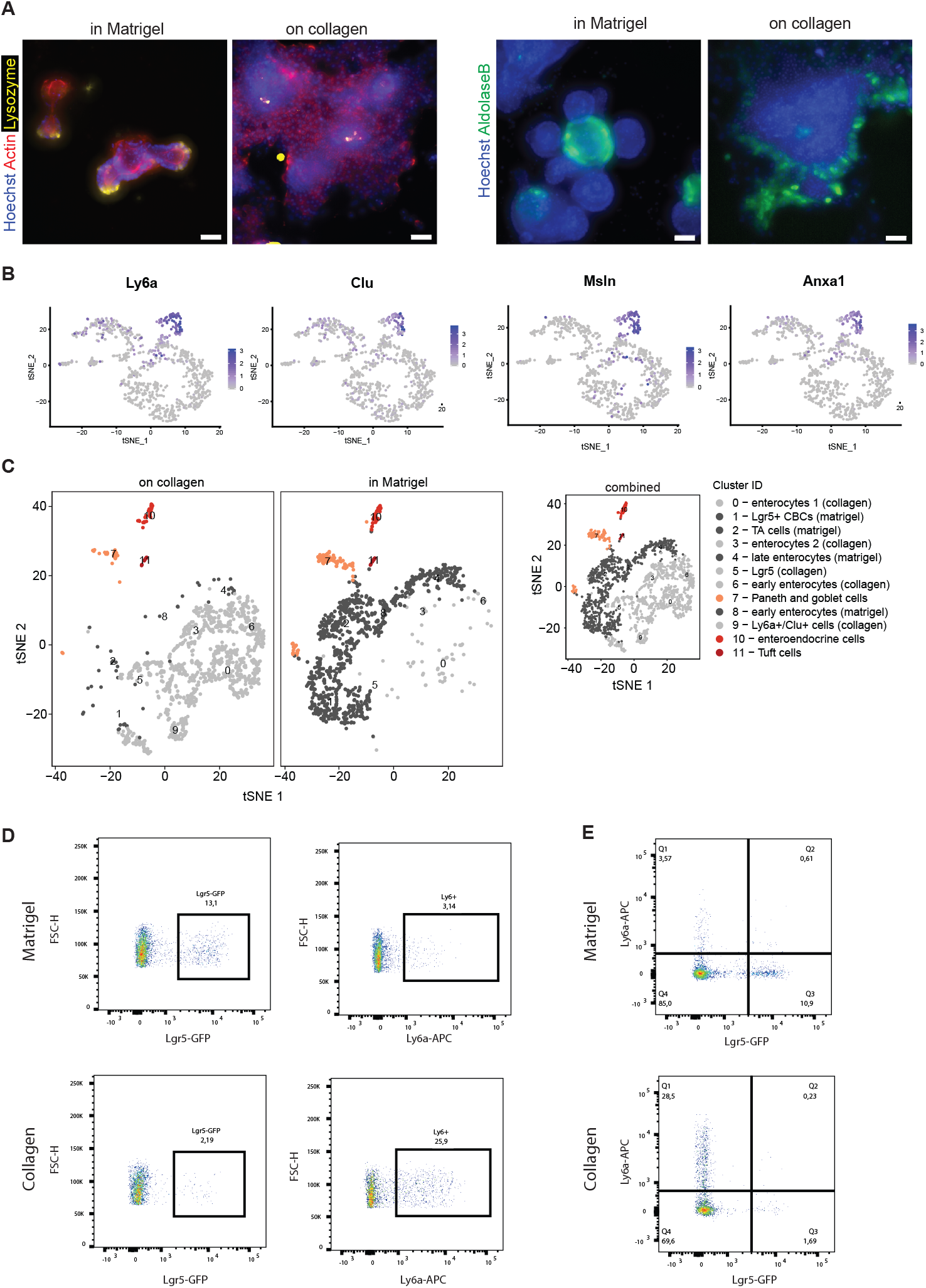
**a**, Differentiated Paneth cells (Lysozyme+) and enterocytes (Aldolase B+) are present in Matrigel as well as in collagen cultures. Scale bar = 100um. **b**, Cells expressing fetal-like genes (Ly6a, Clu, Msln, Anxa1) are highly enriched in cluster 7. **c**, Clustering analysis of cells from Matrigel and collagen combined (small insert). For visualisation, tSNE plot is split by origin of cells, either Matrigel or collagen. Dark grey points are nearly exclusively derived from Matrigel, and light grey dots are nearly exclusively derived from collagen. Secretory cells (orange-red) cluster together indicating a similar transcriptional profile irrespective of the matrix they are grown in. **d**, Representative flow cytometry for Lgr5-GFP and LY6A-APC on small intestinal cultures from Matrigel and collagen (n = 2). **e**, There were no cells stained positive for LY6A that also expressed Lgr5-GFP (Q2) indicating independent cell lineages.

**Fig. S4.**
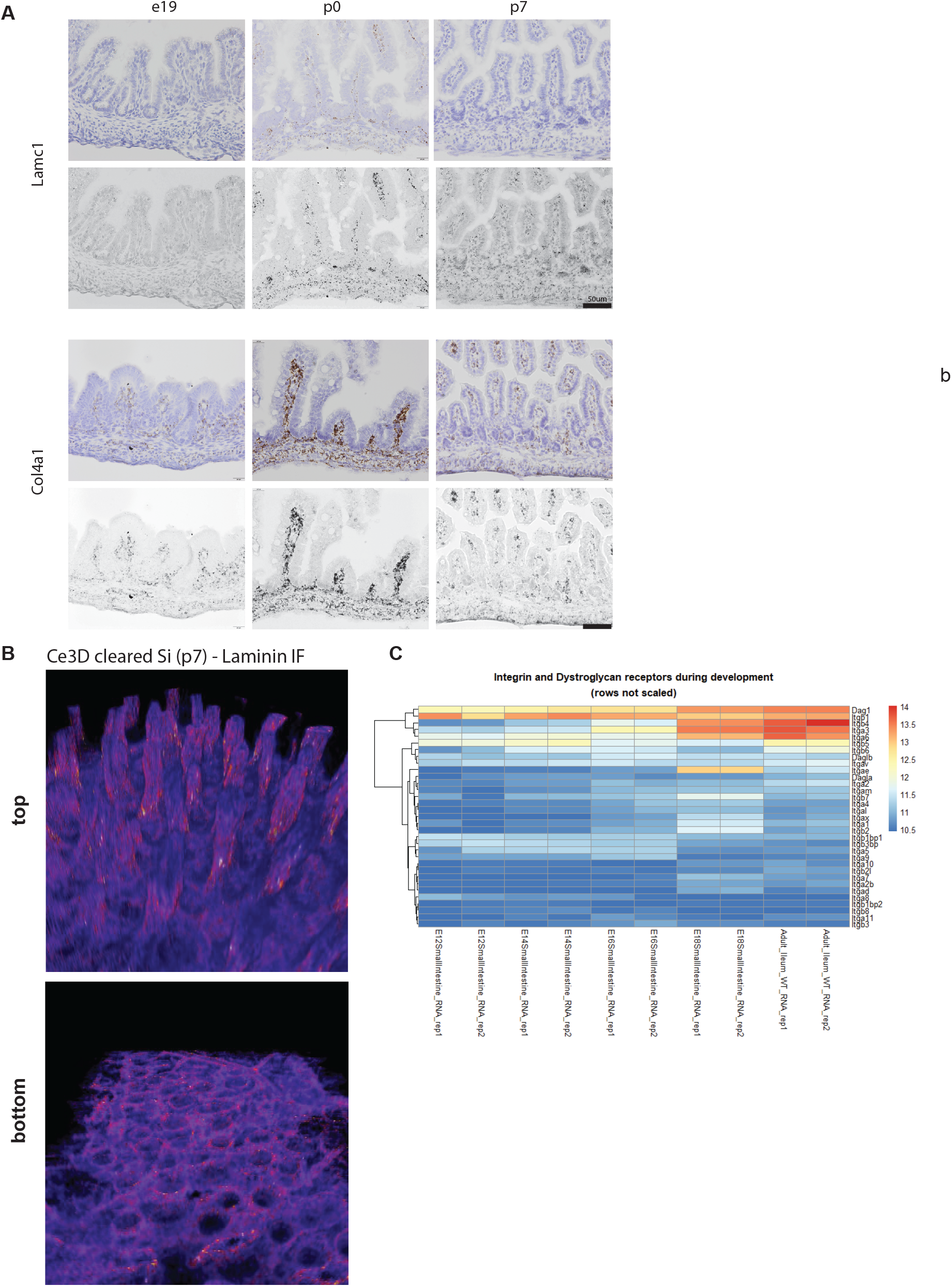
**a**, RNA in situ (RNAscope). RNA in situ hybridization for Laminin C1 and Col4a1 shows highest expression at birth (p0 - as shown in Fig. 3h) and is nearly exclusively found in the mesenchyme. Scale bar = 50um. **b**, Cleared tissue (with Ce3D protocol) from small intestine 7 days after birth (P7) shows underlying laminin network, in the villi (top) and surrounding nascent crypts (bottom). **c**, RNAseq analysis of small intestinal epithelial cells during development (GSE115541) (24) shows expression of genes encoding Integrins and Dystroglycans. Top cluster of genes shows change in expression between E12-E16 and E18-Adult and is highlighted in Figure 4A.

